# CycleGRN: Inferring Gene Regulatory Networks from Cyclic Flow Dynamics in Single-Cell RNA-seq

**DOI:** 10.1101/2025.11.12.688126

**Authors:** Wenjun Zhao, Elana J. Fertig, Genevieve Stein-O’Brien

**Affiliations:** Wake Forest University; Institute for Genome Sciences, University of Maryland Baltimore; Johns Hopkins University

## Abstract

Oscillatory processes such as the cell cycle play critical roles in cell fate determination and disease development, yet existing gene regulatory network (GRN) inference methods often fail to account for their dynamic nature. We propose *CycleGRN*, a novel framework that treats cell cycle gene expression observations as an invariant measure of a stochastic differential equation and learns from data a dynamical system that fits cycling biological processes. Using a directed graph constructed along the inferred flow field in the cell space, we estimate Lie derivatives for all genes, enabling velocity inference beyond the cell cycle subspace. To quantify regulatory interactions, we introduce a time-lagged correlation operator between any pair of genes supported on the flow-aligned directed graph, which respects the intrinsic geometry of the data manifold and allows temporal ordering consistent with the underlying oscillatory process. The method requires only raw gene expression data at single-cell resolution and a list of cycle genes, without temporal binning or splicing dynamics. We evaluate our method on four synthetic datasets generated from mechanistic models with known network structures with oscillatory subnetworks, and on a mouse retinal progenitor single-cell RNA-seq dataset spanning three cell types and a knockout condition. Across all settings, our method consistently ranks among the top-performing approaches and demonstrates strong recovery of oscillatory and directional interactions.

## 1 Introduction

Oscillatory processes such as the cell cycle play a central role in cell fate determination, tissue homeostasis, and disease progression, including cancer and developmental disorders [17, 27, 32]. Beyond cell cycle states, even individual pathways, such as *Notch* signaling [15], can demonstrate oscillatory dynamics. These processes induce structured temporal patterns in gene expression, often reflecting periodic or quasi-periodic dynamics. While such oscillations are readily detectable at the population level, they are typically only indirectly captured in single-cell RNA-sequencing experiments, where each cell is measured at a single time point and thus requires additional methods to infer its developmental or phase state [25].

Despite their biological importance, oscillatory processes have not been fully leveraged in current gene regulatory network (GRN) inference methods. For instance, a common practice in single-cell analysis is to treat cell cycle effects as unwanted confounders [6, 21], removing or regressing out cell cycle–associated genes prior to downstream analyses such as trajectory inference. While this can reduce technical variability, it also discards valuable information about intrinsic temporal structure that could inform regulatory relationships.

We hypothesize that by constraining GRN inference using an estimated invariant oscillatory process, we can better recover the temporal ordering of regulatory events and improve causal inference in oscillatory systems. Intuitively, the invariant process acts as a latent clock shared across cells, capturing the periodic structure underlying gene expression variation. A natural approach is to leverage trajectory inference methods to learn a pseudotime for cell cycle [10]. For instance, Tricycle [31] identifies a template with clear cyclic structure, and uses transfer learning trained on large real datasets to map new datasets onto this template. Empirical evidence using such pseudotime showed improvement in accuracy [30], but required dividing data into discrete bins, which may fail to capture the continuous and periodic nature of the cell cycle.

To address this limitation, we propose to learn the latent cyclic dynamics directly from the data, without inferring explicit pseudotime. Intuitively, cycling genes trace out an orbit in expression space; we seek a vector field whose invariant distribution matches the observed point cloud of these genes [5, 28]. The learned flow defines a local velocity field that respects the cyclic structure of the underlying process. We then extend this invariant flow to all genes by computing cycle-aware directional derivatives, which estimate gene-wise rates of change along the learned flow using a flow-aligned transition operator.

Finally, we define a family of time-lagged gene-gene correlation operators built on the graph that follows the invariant flow of cells, enabling the estimation of directional dependencies between genes without explicit temporal binning or splicing information. This results in a signed, asymmetric gene–gene correlation matrix that reflects potential causal ordering along the oscillatory process. Figure 1 provides a schematic overview of the *CycleGRN* framework. We demonstrate the performance of our method on four simulated datasets with specified oscillatory subnetworks from HARISSA [26], and three real datasets of different cell types with active cell cycles [7], showing superior performance across all tasks, compared to traditional methods that do not leverage knowledge on the oscillatory process.

**Figure 1.**
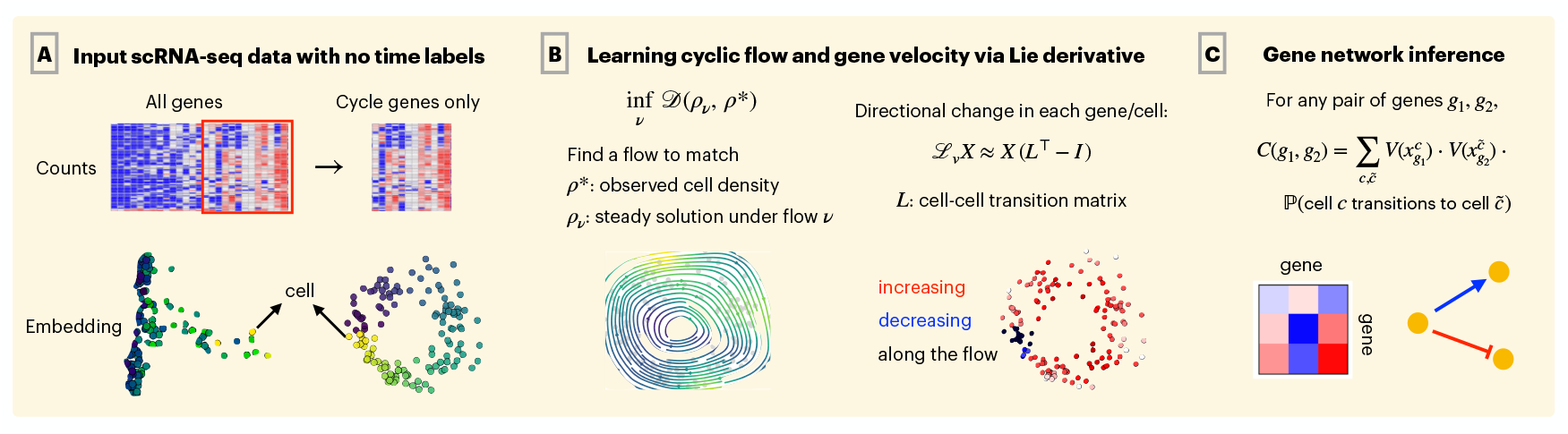
Learning dynamic gene regulation with CycleGRN. (A) Input: scRNA-seq data and a list of cycle genes. (B) A continuous flow in the subspace of cycle genes is learned on this embedding to reproduce the observed cell density. Each gene’s directional change along the flow is estimated. (C) Comparing these directional changes across cell transitions yields time-lagged correlations that reconstruct directed regulatory interactions.

### Our contributions

(1) We introduce a method that learns invariant cyclic dynamics directly from scRNA-seq data via PDE-constrained optimization, requiring neither time labels nor splicing information. (2) We develop a flow-aligned Lie derivative that extends the learned dynamics from cycle genes to the full transcriptome, yielding gene- and cell-specific velocity estimates. (3) We define a family of flow-aligned time-lagged correlation operators that exploit the learned dynamics to infer directional and signed gene regulatory interactions.

### Scope and identifiability

CycleGRN is designed for settings in which the dominant latent dynamics form a one-dimensional cyclic manifold (topologically *S*^1^), such as the cell cycle or other recurrent biological programs. In this regime, the invariant distribution determines the underlying flow up to a global orientation and time reparameterization. Our objective is therefore not to recover absolute biological time, but to infer flow-consistent directional influence within this identifiable equivalence class. Resolving the global orientation requires minimal external information (e.g., canonical phase markers), serving as an identifiability constraint rather than an imposed regulatory prior. We do not claim applicability to arbitrary non-cyclic or multi-scale dynamics.

## 2 Methodology

Given a single-cell gene expression matrix *X* ∈ ℝ^*G*×*C*^, where rows correspond to genes and columns to cells, we partition the gene set into two disjoint subsets, *G* = *G*^inv^ ∪ *G*^other^, where

- *G*^inv^: genes associated with an invariant process (e.g., cell cycle or Notch signaling);
- *G*^other^: all remaining genes reflecting other dynamics (e.g., differentiation or reprogramming).

Let 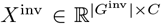 denote the submatrix of *X* restricted to *G*^inv^. The ultimate goal is to use the invariant dynamics in the space of *G*^inv^ to constrain the temporal order of regulatory events, and infer causality that respects the cell cycle – for the full set of genes including both cell cycle genes *G*^inv^ and *G*^other^. We describe the pipeline to achieve this in the following subsections.

### 2.1 Learning the dynamics of cell cycle genes

In single-cell datasets, it has been shown that sampling a population of unsynchronized cells allows the dataset to effectively capture temporal dynamics. Numerous trajectory inference methods and RNA-velocity methods have been developed to infer these dynamics [20], and shown to be successful at uncovering trajectories associated with cell fate decisions. However, these methods often rely on knowing a pre-defined initial state and are fundamentally manifold dependent, limiting the ability to infer more complex dynamics in cyclic processes that lack a unidirectional trajectory.

In the context of chaotic dynamical systems, it was first proposed by Yang et al [5, 28] that even without knowing the actual time label, one can still learn from data the dynamical system by matching the observed density and the invariant measure of a flow. Given a noisy point cloud representing the expression of the set of genes in *G*^inv^, denoted as 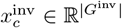 with *c* = 1, …, *C* for *C* cells in total, we represent the distribution of samples as a histogram over predefined bins of expression values. For each bin *B*, we count how many samples fall inside it and normalize by the total number of cells:

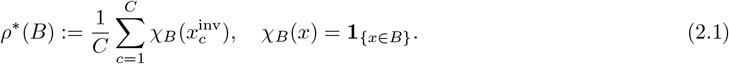

We aim to construct a flow *ν*_*θ*_(*x*^inv^) where *θ* ∈ Θ denotes the parameters in a simple neural network parameterizing the flow in the cell space. The optimal flow is solved through the optimization:

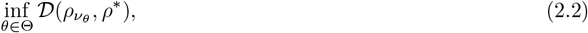

where *ρ*^*∗*^ is the density obtained from single cells representing cell cycle genes, 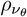 is the steady-state density induced by the flow *ν*_*θ*_, and *𝒟* is a metric that quantifies the discrepancy between two distributions, such as the quadratic Wasserstein distance and Kullback–Leibler divergence. At optima, the steady-state density 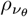 best matches the observed distribution of *ρ*^*∗*^.

We next describe how 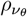 is connected to our parameterized flow *ν*_*θ*_. Derived from the Fokker–Planck equation of a stochastic differential equation 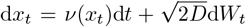, the steady-state solution *ρ*_*ν*_ (*x*) induced by the flow *ν*(*x*) satisfies:

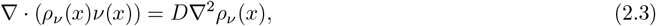

where *ρ*_*ν*_ (*x*) represents the density that is invariant under flow *ν*(*x*), and *D* is a positive constant representing diffusion scale. Following the approach by [5], we solve the equation using a finite volume numerical solver, compute the gradient of the objective based on the adjoint-state method and backpropagation for the parameters in *ν*(*x*), and use gradient-based optimization algorithms to iteratively update them when solving (2.2).

#### Remark on existence and identifiability

The existence of a solution to (2.2) depends on the parameterization class of *ν* [5]. When *ν* is parameterized by a smooth (i.e. *C*^*∞*^) neural network, a regular solution to the steady-state Fokker–Planck equation exists under mild conditions. In practice, we use a two-layer fully connected neural network with tanh activation throughout all experiments.

Beyond existence, the learned flow is not uniquely identifiable from the invariant measure alone. When the underlying dynamics lie on a one-dimensional cyclic manifold (topologically *S*^1^), the invariant density determines the flow up to (i) global orientation reversal and (ii) time reparameterization along the cycle. Thus, additional information is required to fix the direction of traversal.

In our setting, this ambiguity is resolved using minimal biological orientation signals (e.g., canonical phase markers such as S/G2M genes [21]). This does not impose edge-level regulatory priors, but serves as an identifiability constraint analogous to selecting a temporal root in pseudotime methods. A detailed discussion of identifiability under cyclic dynamics is provided in Supplementary Section A.

### 2.2 Cycle-aware Lie derivatives of all genes

The learned flow *ν*(*x*) is defined only on the cycle subspace and therefore must be lifted to the full *G*-dimensional gene space. We approximate this lifting by constructing a cell-based operator that acts on expression profiles and captures directional changes along the flow field. Conceptually, this corresponds to computing a discrete Lie derivative of each gene’s expression with respect to the flow *ν*.

We first construct a transition matrix 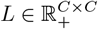 representing a directed *K*-nearest neighbor graph [3, 8]. Each entry *L*(*i, j*) measures how likely cell *i* is to transition toward cell *j* following the local flow direction 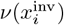:

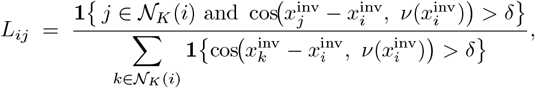

where cos(*a, b*) denotes the cosine similarity, and *δ ≥* 0 is a tolerance threshold. This construction ensures that *L*_*ij*_ is nonzero only for neighbors *j* whose displacement vectors are well aligned with the local flow direction 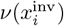. The denominator normalizes rows of *L* to sum up to 1.

Given this directed adjacency matrix, the velocity of other genes can be estimated by a discrete forward difference:

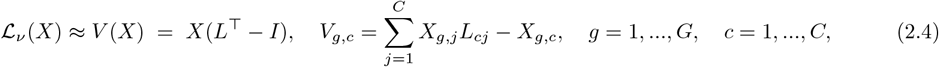

where *X* ∈ ℝ^*G*×*C*^ is the full gene-by-cell matrix. Multiplying *X* by *L*^⊤^ propagates expression values one step forward along the directed cell-cell graph, analogous to a Lie derivative along the flow field, in the operator-theoretic view [8, 19]. The term *X*(*L*^⊤^ *− I*) thus represents the expected forward change of each gene’s expression across cells. Each element *V*_*g,c*_ quantifies the rate of change of gene *g* in cell *c*.

#### Remark on comparison to other velocity methods

Most existing velocity methods, such as RNA velocity [4, 12], rely on spliced and unspliced read counts, which may not be available when sequencing depth is limited. Other approaches incorporate additional information, for example, gene regulatory interactions [13], or rely on binning cells into experimental or pseudotime intervals [29]. In contrast, our method does not require splicing dynamics or explicit temporal binning. It operates solely on a predefined list of cell cycle genes to construct a flow field, and propagates this information to all genes while preserving the continuous granularity of cellular dynamics without discretizing time, under the assumption that the underlying dynamics are truly cyclic, a notoriously hard case for modern trajectory inference tools such as optimal transport (e.g., commented by Supplementary Material of Klein et al [11], “On the limitations of optimal transport for recovering cell cycles”). A comparison between flows found by optimal transport and our method can be found Figure 3, showing that optimal transport may still fail to identify a coherent cyclic flow even with suitable temporal binning.

### 2.3 Time-lagged correlation on the directed graph

The idea of approximating the manifold through a transition matrix can be extended beyond a single step to capture multi-step propagation along the velocity field *ν*. To propagate information along the local flow, we construct a lagged neighborhood operator built from *L*. Given a scalar *α* ∈ [0, 1), we define the propagation matrix

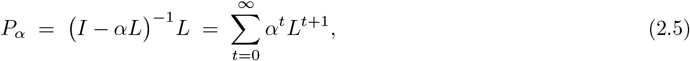

where the latter expansion as a Neumann series corresponds to repeatedly applying the local flow-aligned transition matrix with exponentially decaying weights *α* raised to the power of *t*. When *α* = 0, the propagation operator reduces to *P*_0_ = *L*, which corresponds to a single-step update along the directed neighborhood graph. Because each row of *L* sums to one, this is equivalent to applying one step of a Markov chain whose transition probabilities are supported on the *K*-nearest neighbors aligned with the local flow direction.

We then follow the approach of [29] to define a generalized notion of time-lagged correlation using the propagation matrix *P*_*α*_, after normalizing each gene velocity *V*_*g*_ to have unit variance. For any pair of genes (*g*_1_, *g*_2_), we define

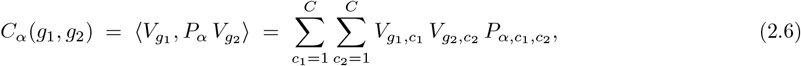

where *V*_*g,c*_ denotes the estimated velocity of gene *g* in cell *c* as in Equation (2.4). The resulting matrix *C*_*α*_ ∈ ℝ^*G×G*^ thus quantifies the empirical correlation between gene-wise rates of change across cell pairs that are likely to be related as ancestor and descendant along the cycle.

#### Remark on the properties of *C*_*α*_

The matrix *C*_*α*_ is signed: positive values indicate that two genes tend to change in the same direction along the flow, whereas negative values indicate opposing changes. Moreover, because *P*_*α*_ encodes time-lagged propagation, *C*_*α*_ is generally asymmetric, allowing one to infer potential directional relationships. Specifically, if changes in *g*_1_ consistently precede those in *g*_2_, then *C*_*α*_(*g*_1_, *g*_2_) will differ from *C*_*α*_(*g*_2_, *g*_1_), suggesting a source–target (causal) ordering between the two.

## 3 Results

We benchmark our algorithm on two types of datasets: (1) synthetic datasets simulated from mechanistic models [26], and (2) real mouse retinal progenitor cells undergoing cell cycle transitions during development [7] in three cell types and one knockout condition. We compare against the following representative algorithms:

- HARISSA and CARDAMOM [26]: mechanistic models for RNA count data that infer regulatory networks based on discretized temporal bins and expression matrices. CARDAMOM extends HARISSA by better exploring the energy landscape and metastable states.
- GENIE3 [1] and GRNBoost2 [14]: decision tree–based algorithms that do not use temporal information. These were ranked among the top-performing methods by BEELINE [18] for both curated and real datasets, with GRNBoost2 serving as a computationally efficient implementation of GENIE3.
- SINCERITIES [16]: a time-aware algorithm that quantifies Granger causality through a linear regression model on Kolmogorov–Smirnov statistic between consecutive time pairs.

For those methods that require time coordinates, we use either the Tricycle coordinates generated by [31], which use transfer learning to project cell cycle genes onto a common periodic template, or the simulation/experimental times when available. Performance is quantified by the area under precision curve (AUPR) and its variants.

Because the current implementation of the finite-volume solver for the Fokker–Planck equation is limited to two dimensions, in our proposed method *CycleGRN*, we learn the flow *ν* based on the first two principal components of cycle genes for synthetic data. For real datasets, we report results using both the first two principal components and the two-dimensional embedding produced by Tricycle [31]. Gene velocities are then computed using Equation (2.4) and normalized to unit variance across cells prior to computing time-lagged correlations. The lag parameter *α* in Equation (2.5) is set to 0. More details can be found in Supplementary Information Section 1.

### 3.1 Results on simulations

We first evaluate the performance of our method on simulated datasets where the ground truth regulatory network is known and can be directly compared with the inferred results. HARISSA [26] is a software framework based on a mechanistic model that simulates RNA counts given any user-specified network. We used three benchmark networks: (1) FN4, a 4-gene network with a feedback loop; (2) CN5, a 5-gene network exhibiting cyclic dynamics; and (3) FN8, an 8-gene network with a feedback loop. To further mimic the complexity of real signaling pathways, we also simulated additional datasets inspired by the *Notch* pathway, including the motif between *Notch1, Dll1*, and *Hes1*. For each network, we generated 10 replicates with different random seeds, each containing 1,000 cells. Ground-truth networks and example simulated datasets are shown in the top two rows of Figure 2, with additional visualizations in Supplementary Figure S1.

**Figure 2.**
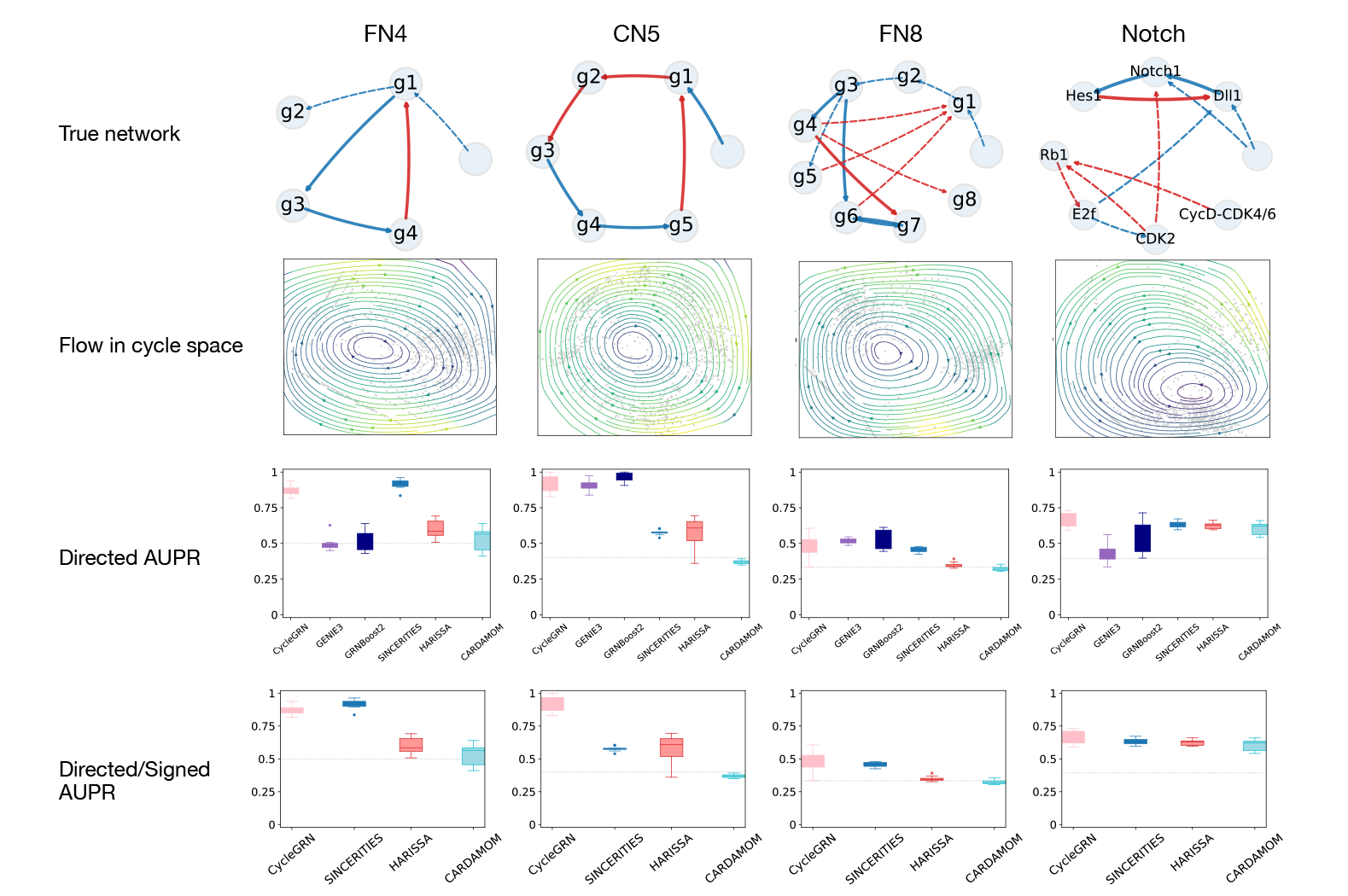
Results on oscillatory networks simulated by HARISSA [26]. Top row: ground-truth gene regulatory networks, where solid lines indicate edges associated with cycle genes used to learn the invariant flow, and edge color represents activation (blue) or repression (red). Second row: invariant flow ν_θ_(x) learned by our algorithm on the oscillatory subspace. Third row: area under the precision–recall curve (AUPR) for directed regulation. Bottom row: signed AUPR that additionally accounts for the regulation type (activation vs. inhibition). GENIE3 and GRNBoost2 are omitted because they do not predict regulation type.

**Figure 3.**
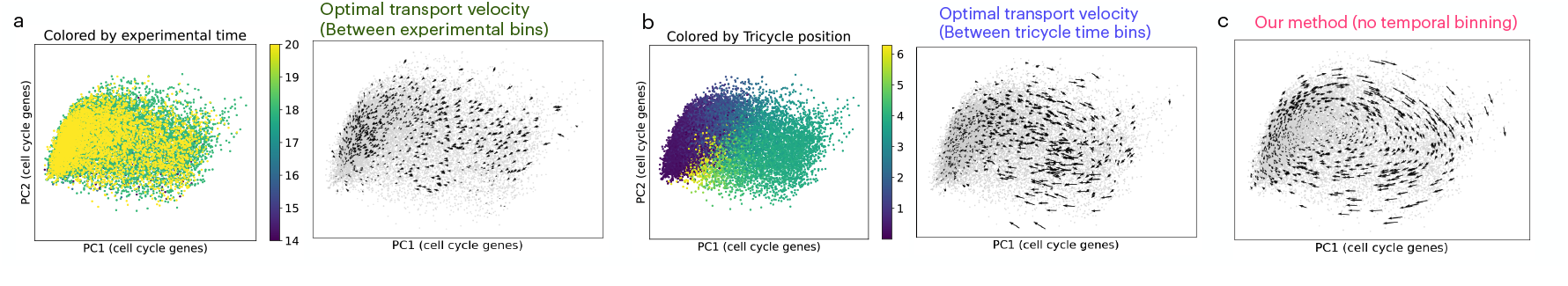
Comparison of velocity fields on late RPCs learned using (a) OT velocity [29] using experimental time, (b) OT velocity [29] using Tricycle phase divided into 20 equally-sized bins, and (c) our invariant dynamics without any time labels. Optimal-transport–based velocity fields fail to recover a coherent cyclic flow under either temporal coordinate, whereas our method recovers a consistent circular dynamics.

Each dataset also records the simulated time point for every cell, visualized by color in Supplementary Figure S1. To mimic experimental conditions, we intentionally introduced long lags between consecutive time points so that simulated “experimental time” does not align perfectly with the internal cell-cycle clock. These simulated times were provided to algorithms that require temporal information, including SINCERITIES [16], HARISSA, and CARDAMOM [26]. HARISSA and CARDAMOM both infer networks by directly fitting the mechanistic model used to generate the data, with CARDAMOM offering better exploration of the energy landscape, metastability, and computational efficiency. In contrast, our algorithm does not use simulation time but instead leverages prior knowledge of oscillatory genes (last row in Supplementary Figure S1). Orientation was fixed as described in Section 2.1 using minimal biological markers (the stimulus gene acting as an activator).

We first learn the flow fields on cycle genes and visualize them in the second row of Figure 2. The learned vector field shows smooth transitions in the space of cycle genes and aligns well with the data distribution. We set *K* = 200 and *δ* = 0.95 to construct the transition matrix *L*, and infer the gene regulatory network using the time-lagged correlation operator described above.

We evaluate accuracy on the directed network using two metrics: (1) the area under the precision–recall curve (AUPR) computed on the absolute values of predicted edges, and (2) a signed AUPR that additionally requires the correct regulation type (activation or inhibition). All metrics are scaled between 0 and 1, with higher values indicating better performance. Results are shown in the bottom two rows of Figure 2; results for undirected prediction are in Supplementary Figure S2. For the two smaller networks, FN4 and CN5, our method consistently ranks top in all metrics compared to any other method. This is not surprising, since the subnetwork used as cycle genes takes a significant portion of the original true network and almost dominates the dynamics. For the two larger and more complex networks, FN8 and Notch, we show comparable or marginally better performance even compared to HARISSA and CARDAMOM, which have access to the original mechanistic model used for simulating the data. We also note that all the metrics are compared directly against the ground truth network, which is directed (distinguishes source and target genes) and signed (distinguishes activation and inhibition), demonstrating that our method can infer the temporal information even not used as input of the algorithm. This is particularly advantageous in real single-cell datasets, which typically contain only one or a few asynchronous snapshots that do not align with biological time.

### 3.2 Results on mouse retinal progenitor cells

We further demonstrate the performance of our algorithm on real datasets of mouse retinal progenitor cells (RPCs) [7]. The cells were subset into 27,764 early RPCs, 16,422 late RPCs, and 6,929 neurogenic cells, respectively. We ran the algorithm using 205 cell cycle genes curated from the Gene Ontology term for mouse cell cycle (GO:0007049) [2], and then infer GRNs using 3, 164 highly variable genes. We benchmarked our method against GENIE3 and GRNBoost2 (which do not use temporal information), as well as SINCERITIES with experimental time and with Tricycle positions. Due to the extensive computational cost, we randomly subsampled 1,000 cells for GENIE3 and GRNBoost2 for each cell type. Following BEELINE [18], due to low absolute accuracy, we evaluated performance using the AUPR ratio (AUPR divided by network density) and AUROC.

The cell cycle embeddings are shown in Figure 4 and Supplementary Figure S3, where the point clouds appear as amorphous blobs without clear cyclic structure. Experimental time also misaligns with the cell cycle phase revealed by Tricycle coordinates as shown in Supplementary Figure S3. Though Tricycle coordinates successfully provide an ordering that agrees to known biology, existing method using optimal transport on that time coordinate still fails to recover a coherent cyclic flow as shown in Figure 3. Nevertheless, our method successfully recovers a dynamical system consistent with the Tricycle positions. The recovered flow fields of each cell type are shown in panel (a) of Figure 4. We then compute gene-wise velocities for all genes, where red indicates increasing and blue indicates decreasing expression. Velocities for *Top2a* are shown in panel (b) of Figure 4, peaking at the mid-cycle phase, consistent with rapid transcriptional dynamics during G2 [27], and is markedly reduced in neurogenic RPCs, reflecting their diminished proliferative activity and exit from the cell cycle.

**Figure 4.**
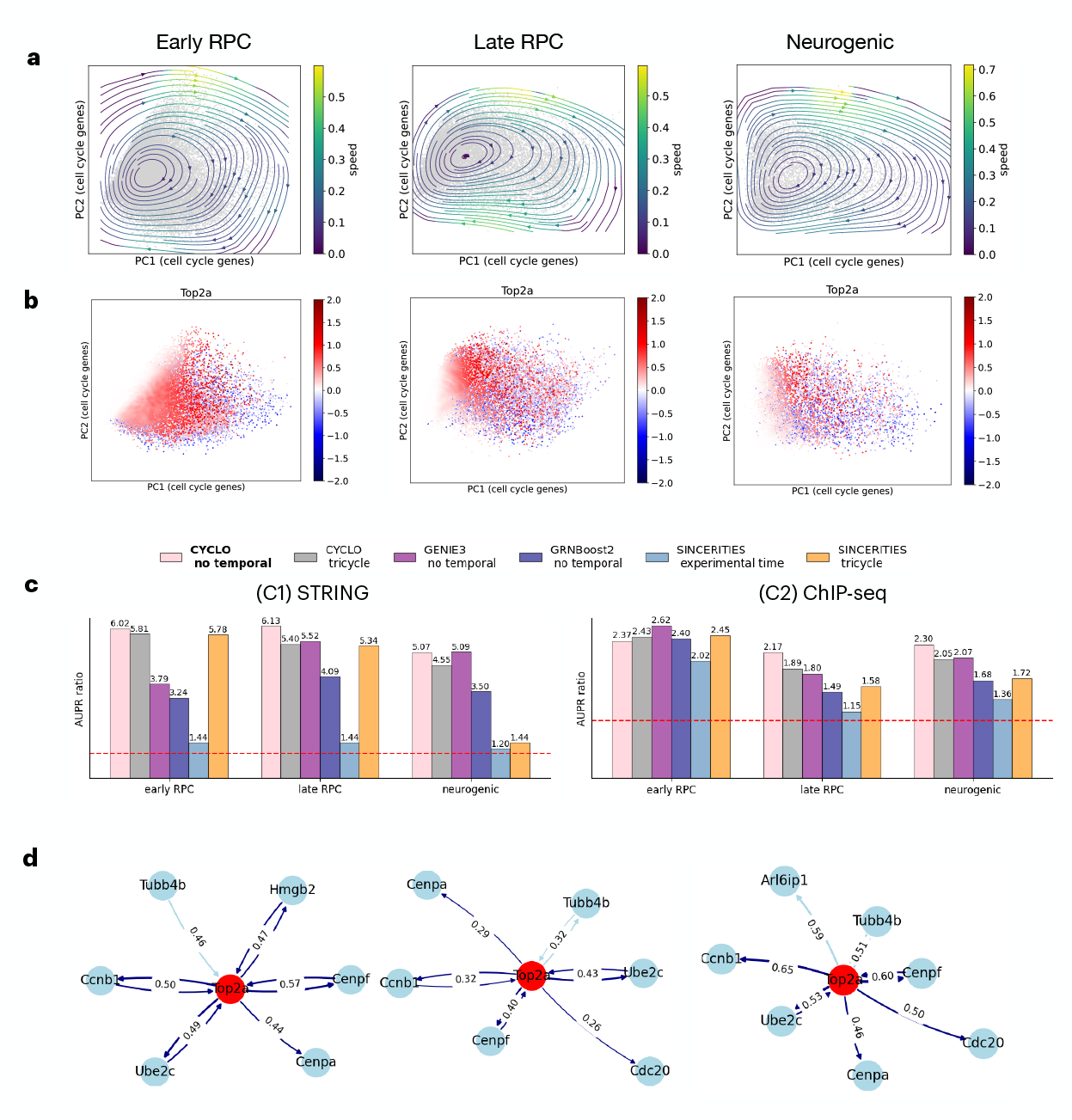
Results on mouse retinal cells in 3 cell types: early retinal progenitor (first column), late retinal progenitor (second column), and neurogenic (third column) cells. Top row: invariant flow learned for each cell type, overlaid on the first two principal components of 205 cell cycle genes. Second row: gene velocity associated with Top2a, estimated by equation 2.4, consistent with the fact that the gene starts to peak in the middle of cycle rather than the beginning. Third row: accuracy of resulting network benchmarked against STRING (b1) and ChIP-seq (b2), quantified by AUPR ratio (AUPR divided by density of network). Our method consistently ranks among the top-performing approaches across all RPC populations, and remains competitive in neurogenic cells where cell-cycle activity is weakened. Bottom row: top 10 highest weighted edges associated with Top2a, with dark blue edges indicating evidence supported by STRING.

Next, we apply the lagged correlation operator to 3,164 genes, selected as transcription factors and pattern markers, to infer directed gene–gene relationships. We benchmark the inferred networks against the STRING database [23], restricted to 80 pattern markers genes curated from non-negative matrix factorization analysis of the complete retinal dataset [7] with CoGAPS [9, 22] within each cell type performed previously on the same dataset. Only regulatory links with at least 80% confidence in STRING were retained for evaluation.

The overall performance, measured by AUPR ratio, is shown in panel (c) of Figure 4 under two complementary benchmarks: (C1) STRING-based evaluation and (C2) ChIP-seq–supported TF–target interactions. Under STRING (C1), our method outperforms all others in early and late RPCs, where cell-cycle activity is pronounced. Under ChIP-seq (C2), performance varies slightly across cell types: GENIE3 shows higher agreement in early RPCs, whereas our method achieves the best or near-best results in late RPC and neurogenic populations. Performance remains comparable to GENIE3 in neurogenic cells, while other methods approach random baseline. Results are robust to the choice of *k* and *τ* (Supplementary Figure S4). Learning the flow directly on the Tricycle embedding yields slightly reduced performance compared to learning on cell-cycle genes [31].

We further examine whether the method improves predictions for cell cycle genes, as hypothesized. For the representative cell cycle gene *Top2a*, we compare the inferred network edges to those predicted by GENIE3 (Supplementary Figure S5), focusing on the top ten edges by weight. GENIE3 correctly predicts 8, 8, and 6 edges across the three cell types, whereas our method predicts 9, 8, and 7, respectively. Notably, our method identified the edge pointing from *Top2a* to *Cenpa*, a G2/M gene that peaks after *Top2a* [27], while GENIE3 missed it. This is likely because this relationship is phase-dependent and directional within the cell-cycle trajectory, and can be blurred by static networks inferred from regression-based methods such as GENIE3. This result is consistent with our hypothesis and highlights the potential of our method for improving regulatory inference in cell types exhibiting strong oscillatory signals.

The methodology is time-lagged correlation–based; therefore, it is crucial to verify whether the inferred networks capture directional causal relationships rather than static associations. To this end, we performed differential GRN inference on (1) regular late retinal progenitor cells (RPCs) and (2) *Nfia/b/x* triple-knockout RPCs, in which the transcriptional regulators *Nfia, Nfib*, and *Nfix* are simultaneously disrupted [7]. This knockout is known to impair the specification of late-born retinal cell types and to cause a loss of Müller glia and bipolar differentiation. The results are summarized in Figure 5.

**Figure 5.**
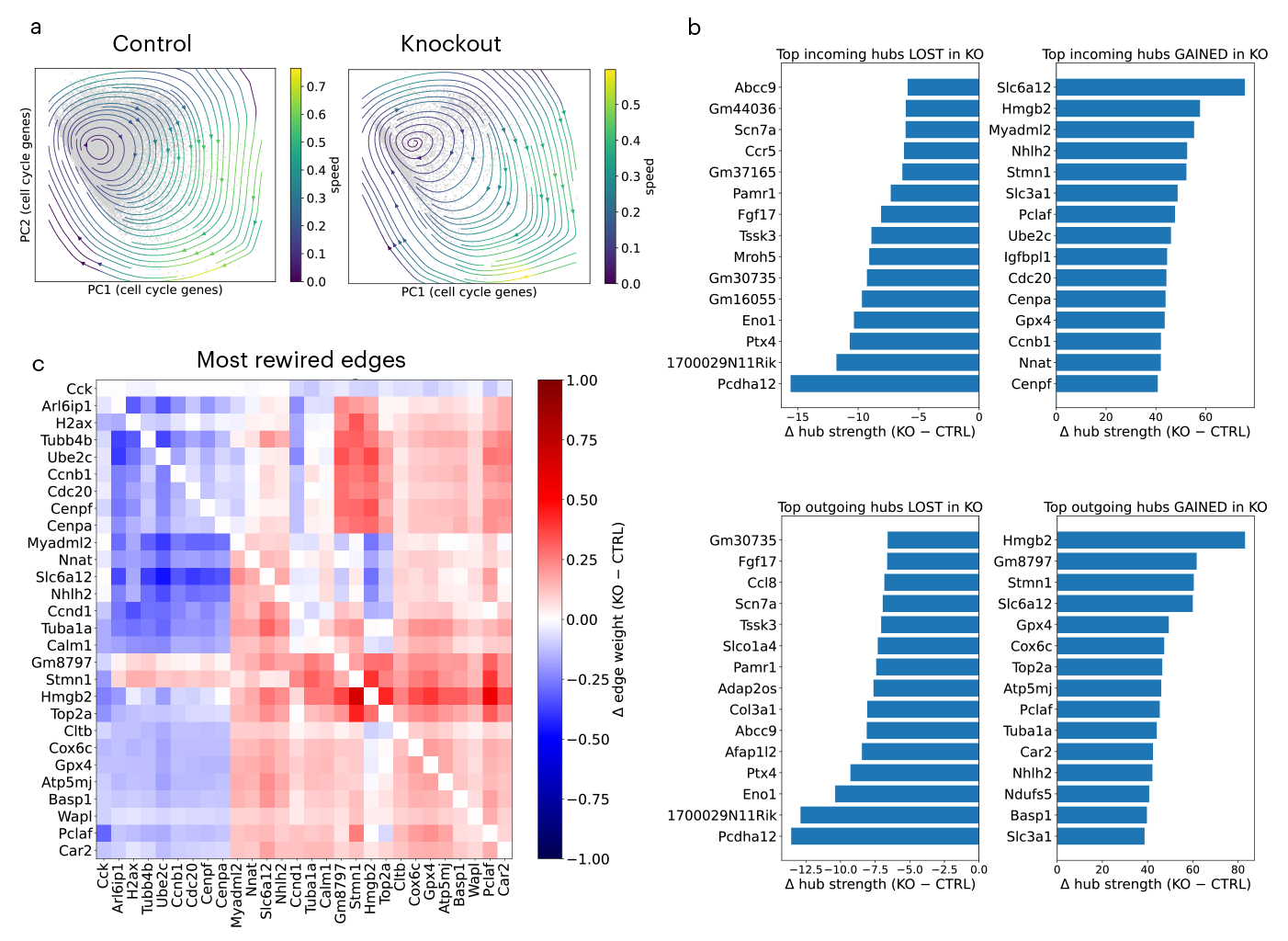
Differential GRN inference on regular late RPCs versus Nfia/b/x triple-knockout RPCs. (a) Vector field learned from cell cycle genes, (b) incoming/outgoing hub genes, and (c) subnetwork for top 50 differential edges.

Figure 5 presents (a) the velocity field learned from cell-cycle genes, (b) the differential incoming and outgoing hub strengths, and (c) the subnetwork formed by the 50 most rewired edges. The inferred network differences recapitulate the experimental phenotype: in the knockout, a reinforced core of cell-cycle regulators (*Hmgb2, Top2a, Ube2c, Cdc20*) gains mutual connectivity and forms a self-sustaining proliferative module, while genes associated with differentiation and late mitotic execution (*Cenpf, Cenpa, Ccnb1, Tubb4b*) lose both incoming and outgoing regulatory edges, reflecting a collapse of coordinated lineage and mitotic control.

This asymmetric rewiring, where upstream proliferative drivers persist while downstream effectors and differentiation targets become decoupled, demonstrates that the inferred network structure reflects directional causality consistent with biological perturbation. The method therefore captures not only known regulatory phenotypes but also distinguishes causal persistence (sustained drivers) from causal disconnection (lost regulation) within perturbed developmental programs.

## 4 Discussion

We have presented *CycleGRN*, a framework that learns an invariant flow in the space of genes corresponding to oscillatory processes such as the cell cycle and leverages it for gene regulatory network inference. The method achieves strong performance on both simulated and real datasets, particularly enhancing inference for cell cycle genes. Although we focus on the cell cycle, the framework naturally extends to other recurrent processes like circadian rhythms and *Notch* oscillations.

There are several avenues for further improvement. First, the current implementation relies on a 2D embedding prior to learning the dynamics. This may fail in settings with substantial cell type heterogeneity, where a simple 2D projection does not reveal a clean cyclic structure. At the same time, the low-dimensional projection can act as a geometric denoising step, stabilizing local distances before flow estimation. Extending the flow learning procedure to higher-dimensional representations could improve robustness across diverse datasets while preserving this regularization effect. Second, as discussed in the methodology, the learned flow is not guaranteed to be unique. Prior work has leveraged time-delay coordinates and Takens’ theorem [5, 24] to address this issue. Although true lagged measurements are not directly available in single-cell experiments, experimental time points or pseudotime estimates could potentially be used to construct suitable surrogate constraints. Third, our algorithm is naturally biased toward oscillatory genes, particularly cell cycle genes, and may not perform optimally for regulatory inference involving other processes. For example, we observed suboptimal recovery of interactions involving *Hes1* (Supplementary Figure S6).

More broadly, biological systems exhibit regulation across multiple temporal and functional scales, from fast cell-cycle oscillations to slower developmental and differentiation programs as well as pathway-specific signaling modules. Capturing these processes within a unified modeling framework remains a key challenge. We envision extending *CycleGRN* toward multi-scale inference, where dynamic regulatory flows corresponding to different timescales and pathways are jointly identified and integrated, bridging cyclic and hierarchical modes of regulation.

## Data and code availability

The HARISSA simulations and scripts to reproduce the experiments are available at https://github.com/WenjunZHAOwO/CycleGRN/tree/main. The mouse retinal dataset is available at https://www.ncbi.nlm.nih.gov/geo/query/acc.cgi?acc=GSE118614.

## A Identifiability of cyclic flows from invariant measure

We characterize the identifiability of latent cyclic dynamics from an observed invariant measure in the presence of diffusion. This analysis clarifies the degree of non-uniqueness in the inferred flow and motivates the additional biological constraints used in the main text.

### Setup

Let the latent cell-cycle coordinate be parameterized by *θ* ∈ *S*^1^ ≃ ℝ*/*(2*π*ℤ). We consider a one-dimensional diffusion process

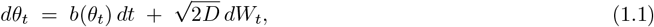

where *b* : *S*^1^ → ℝ is a 2*π*-periodic drift field and *D >* 0 is a constant diffusion coefficient. Suppose the process admits a strictly positive stationary density *ρ*(*θ*) satisfying

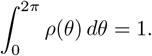

### Stationary Fokker–Planck equation

The stationary density *ρ* satisfies the one-dimensional periodic Fokker– Planck equation

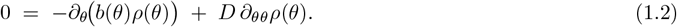

Define the stationary probability current

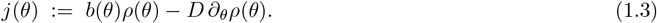

Equation (1.2) implies *∂*_*θ*_*j*(*θ*) = 0, hence

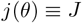

for some constant *J* ∈ ℝ representing the net circulation around the cycle.

### Drift decomposition

Solving (1.3) for the drift yields

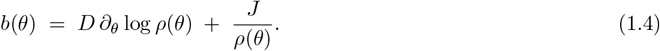

This decomposition separates the drift into a density-determined component and a circulating component corresponding to persistent directional flow around the cycle.

### Identifiability. Proposition

Let *b*_1_ and *b*_2_ be two drift fields on *S*^1^ generating diffusion processes with the same diffusion coefficient *D >* 0 and the same strictly positive stationary density *ρ*. Then there exists a constant *c* ∈ ℝ such that

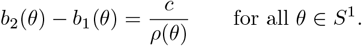

*Proof*. Both drifts admit the representation (1.4) with constants *J*_1_ and *J*_2_, respectively. Subtracting the two expressions gives

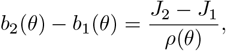

which proves the claim. □

### Interpretation

Thus, on a one-dimensional cyclic manifold, the invariant density *ρ* determines the spatial structure of the drift uniquely, leaving only a single scalar ambiguity corresponding to the magnitude and direction of circulation. This ambiguity cannot be resolved from the stationary distribution alone and reflects a fundamental non-identifiability of cyclic flows from steady-state data. In the main text, we resolve this remaining degree of freedom using biological prior information, specifically known cell-cycle marker genes, which provide temporal orientation along the cycle.

### Remarks

If detailed balance is assumed, then *J* = 0 and the drift is uniquely determined by *ρ* via *b*(*θ*) = *D ∂*_*θ*_ log *ρ*(*θ*). If the diffusion coefficient *D* is unknown, additional non-identifiability arises and cannot be resolved from the invariant density alone.

## B Implementation note on learning the flow

The flow field *ν*_*θ*_ was trained following the implementation of Botvinick-Greenhouse et al [5], without much modification to the original code available at https://github.com/jrbotvinick/Learning-Dynamics-on-Invariant-Measures. The invariant-measure solver and adjoint gradient are implemented using a finite–volume discretization of the Fokker–Planck equation coupled to a neural-network parameterization of the drift field.

### Domain and discretization

We restricted the invariant-gene embedding to a two–dimensional square domain [*−*4, 4] *×* [*−*4, 4] with spatial step Δ*x* = 0.1 and time step Δ*t* = 0.01, yielding *n*_*x*_ = *n*_*y*_ = 81 grid points. The diffusion coefficient was fixed at *D* = 10^*−*3^ and a small teleportation parameter *α* = 10^*−*8^ was added to ensure ergodicity. The empirical steady–state density *ρ*^*∗*^ was estimated from the cell embeddings via a two–dimensional histogram, normalized to integrate to one, and smoothed by a Gaussian kernel with standard deviation *σ* = 2 grid cells. We also note that because the space is discretized and we only use the relative counts of cells within each bin to learn the dynamics, computationally, the method scales very well with number of cells.

### Velocity parameterization

The drift field *ν*_*θ*_(*x*) = (*u*_*θ*_, *v*_*θ*_) was represented by a fully connected neural network with two hidden layers of 100 units each and tanh activation, which guarantees smoothness and consequently existence of a valid solution.

The network was implemented in PyTorch and evaluated on a staggered grid aligned with the finite–volume stencil, enforcing no–flux boundary conditions.

### Optimization and loss

We minimized the Kullback–Leibler divergence between the empirical and model-implied stationary densities,

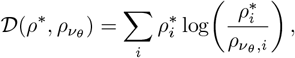

where 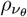 is obtained by solving the steady–state Fokker–Planck equation 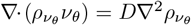. The corresponding adjoint equation was solved analytically in discrete form to compute exact gradients of the loss with respect to *ν*_*θ*_, which were back-propagated through the neural network using a custom `torch.autograd.Function`. Parameters were optimized using Adam with learning rate 10^*−*3^ for 1, 000 iterations, and random seeds were fixed for reproducibility.

## C Supplementary figures

**Figure S1:**
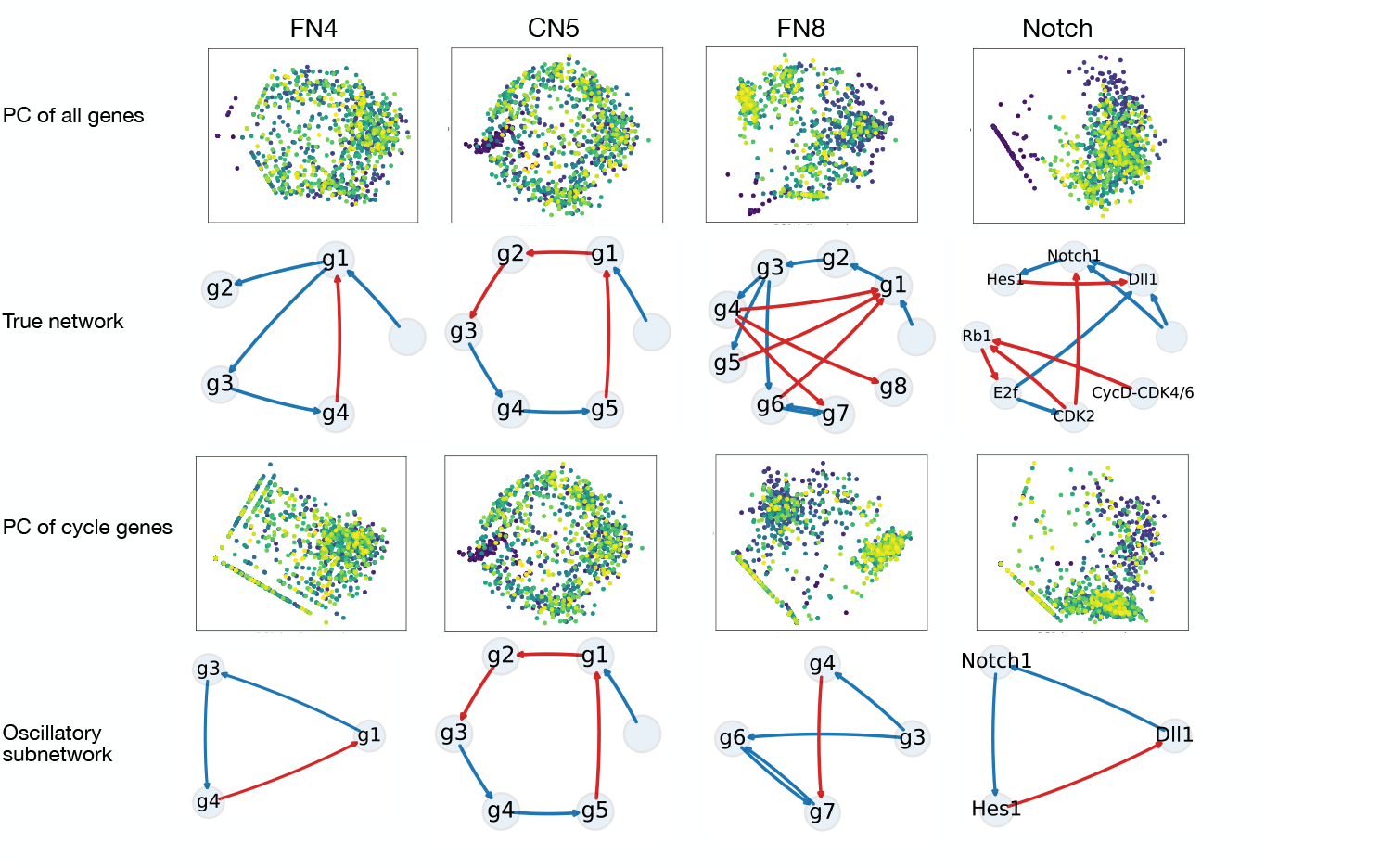
Datasets with oscillatory networks simulated by HARISSA [26]. This figure shows low dimensional visualization and ground truth networks either with all genes (second row) or just a subset that corresponds to oscillatory process (last row). In the principal components, cells are colored according to the time points at which they were simulated.

**Figure S2:**
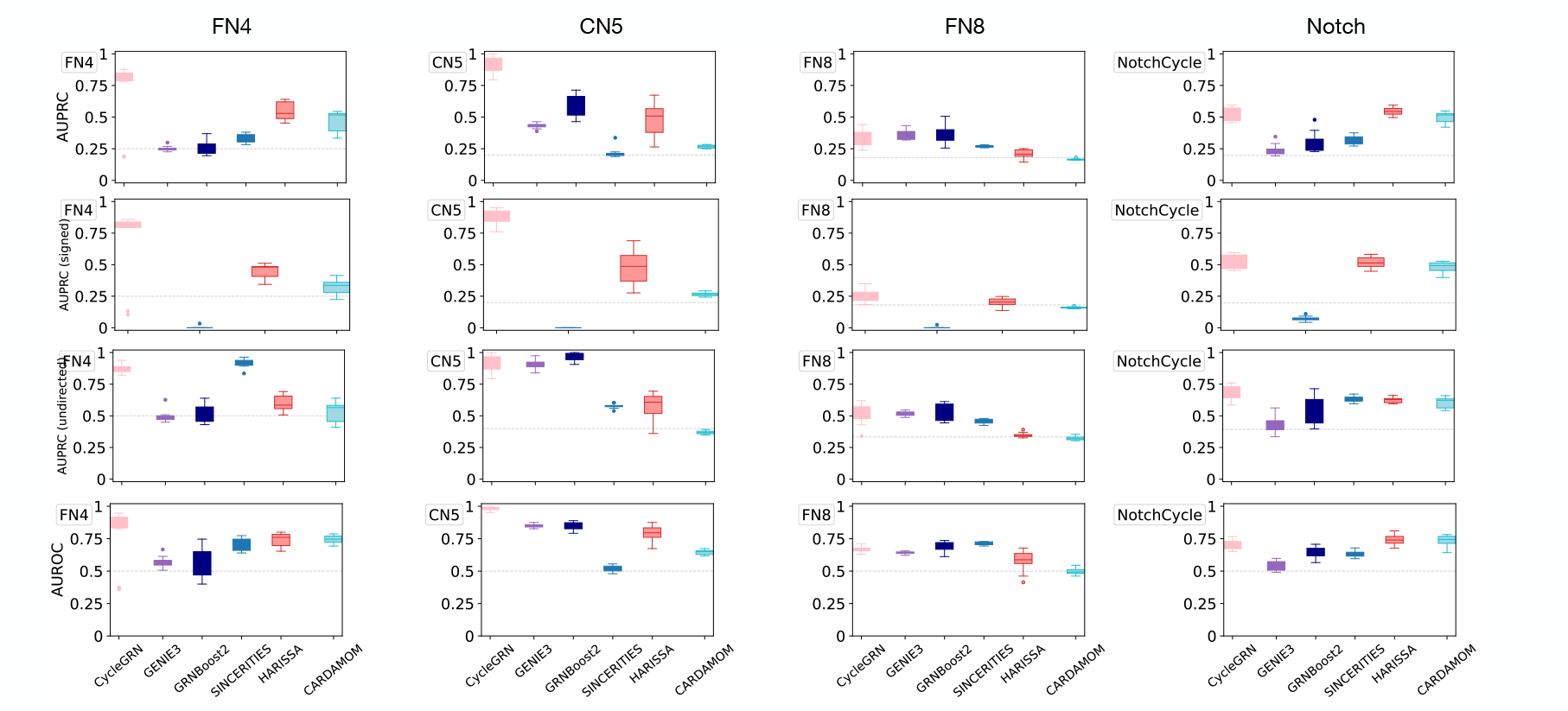
Accuracy metrics on simulated data, resulting from a wide range of algorithms. GENIE3 and GRN-Boost2 do not produce signed prediction and is omitted from the third row on signed AUPRC.

**Figure S3:**
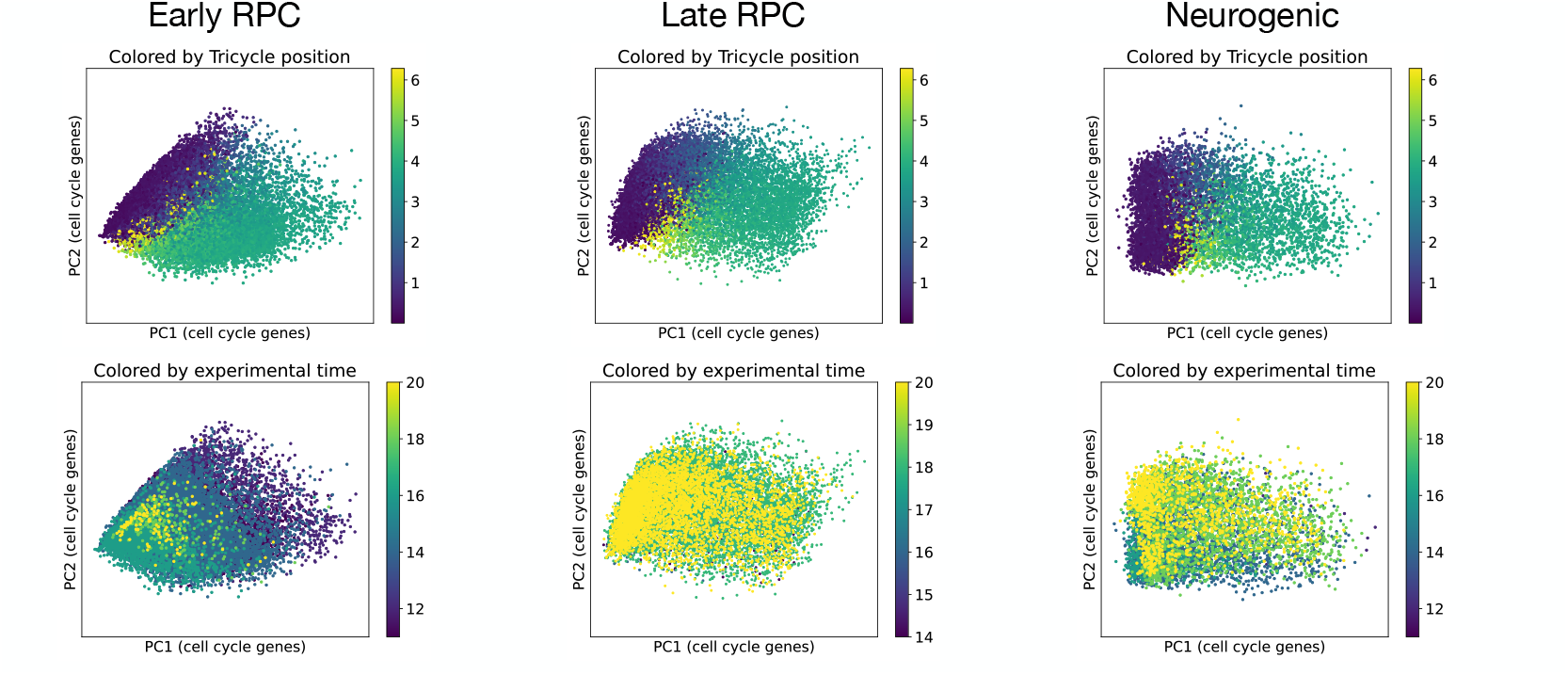
Visualization of mouse retinal cells colored by experimental time (first row) and tricycle coordinates (second row)

**Figure S4:**
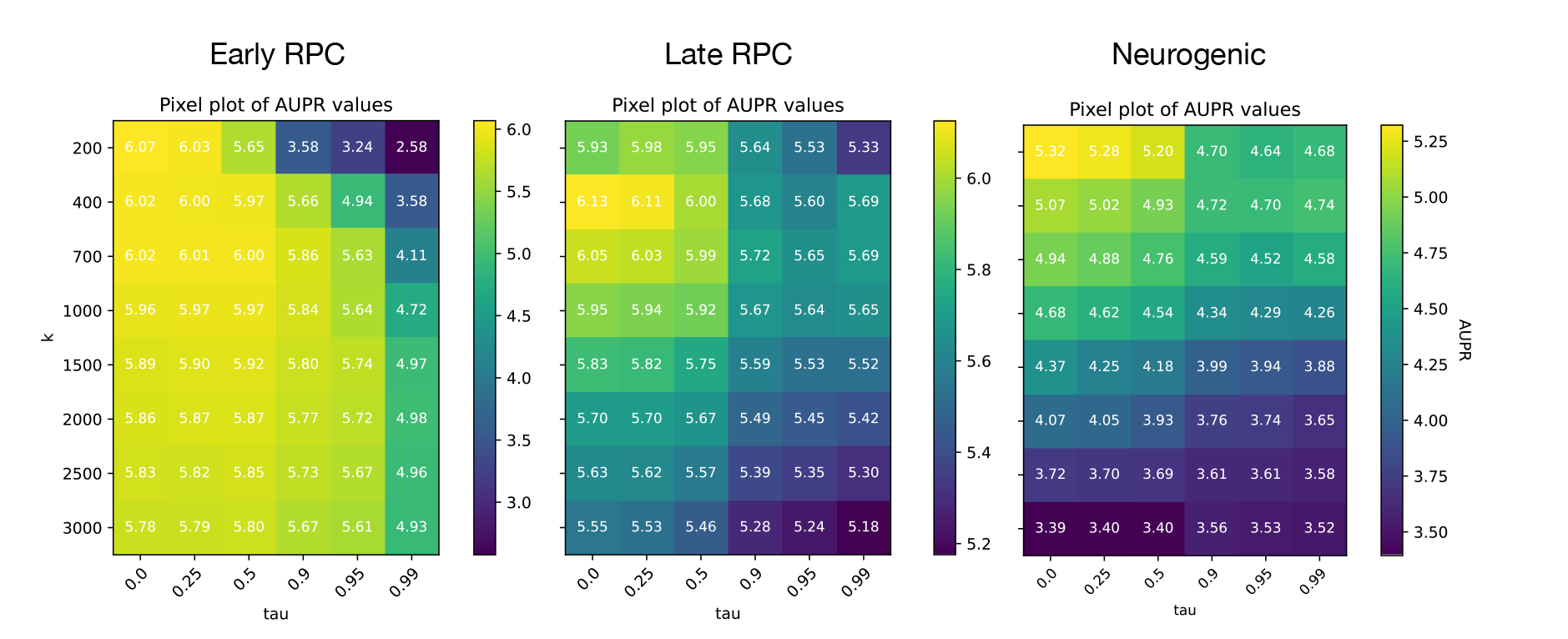
Sensitivity analysis on the hyperparameters τ and k used in constructing the transition matrix on real data. Overall, the performance quantified b y AUPRC ratio i s robust f or m oderate τ a nd k. We have used τ = 0 and k = 400 in all real data studies.

**Figure S5:**
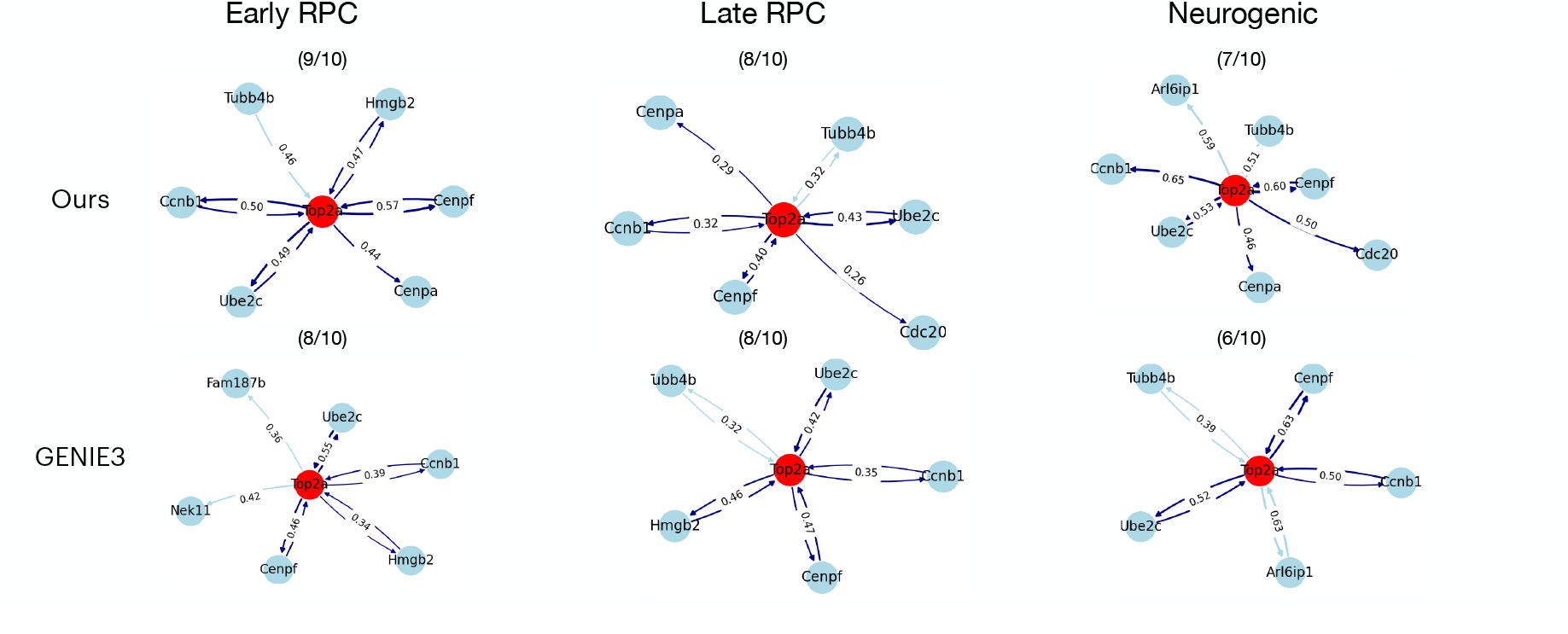
Comparison of networks between GENIE3 and ours on Top2a for top 10 edges. Our method is able to identify one more edge connecting Cenpa, a cell cycle gene that peaks after Top2a, for early RPC and neurogenic cells.

**Figure S6:**
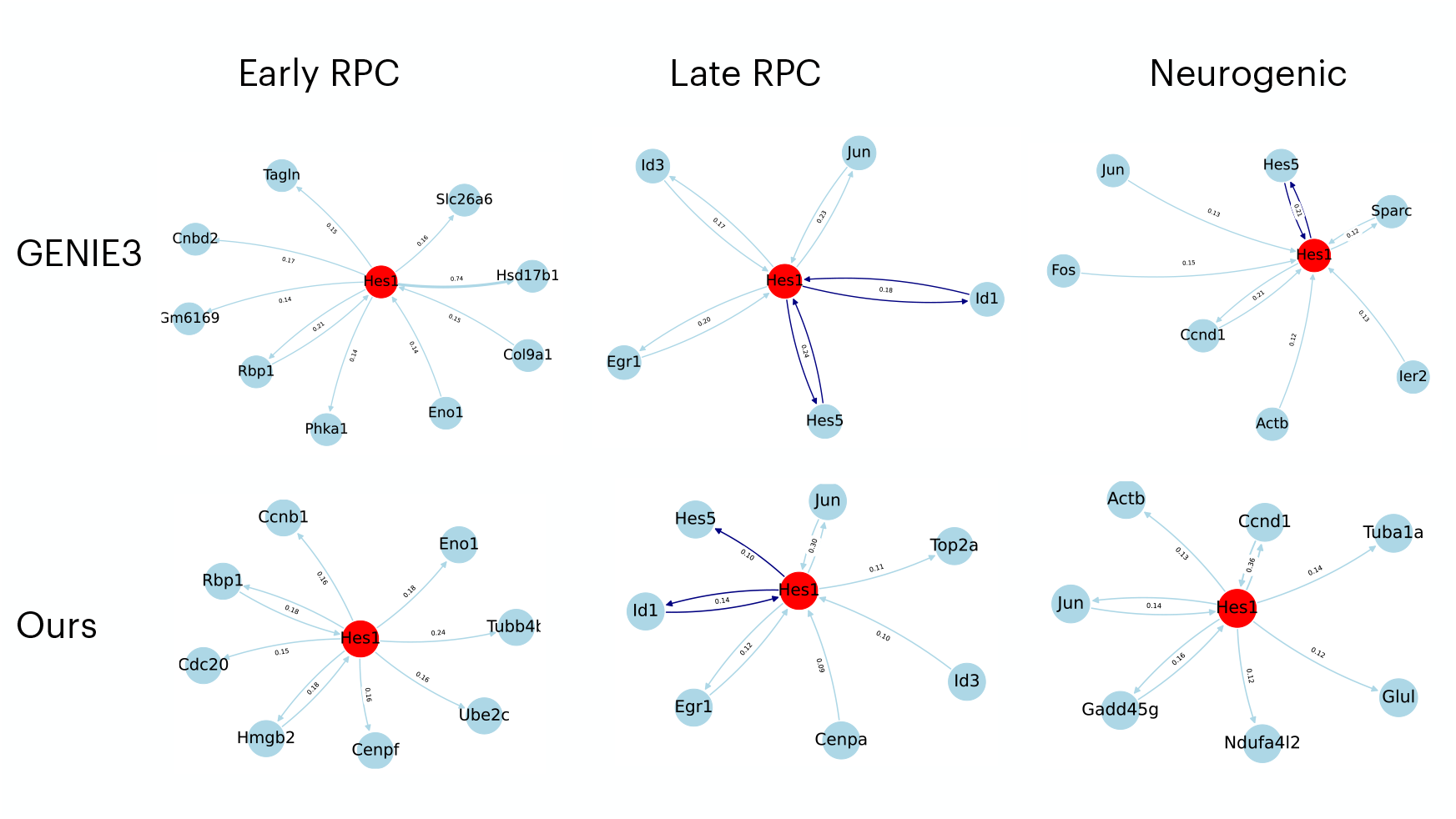
Comparison of networks between GENIE3 and ours on Hes1. As our method prioritized cell cycle genes that do not include Hes1, it can fail to identify edges if its dynamics do not respect the same temporal ordering.

## Footnotes

0 Code with a simple notebook example available at https://github.com/WenjunZHAOwO/CycleGRN/tree/main

